# Intrinsic class C extended-spectrum β-lactamases mediate resistance to oxyimino-cephalosporins in Antarctic *Pseudomonas fluorescens* complex bacteria

**DOI:** 10.64898/2026.02.10.705138

**Authors:** José Coche-Miranda, Iyaray Céspedes-Navarro, Bruno Cardemil, Patricio Arros, Camilo Berríos-Pastén, Ian Pérez, Francisco P. Chávez, Andrés E. Marcoleta

## Abstract

From a One Health perspective, identifying environmental reservoirs of antibiotic resistance determinants with potential clinical relevance is increasingly important. Among resistance mechanisms, β-lactamases are of particular concern because they compromise the efficacy of β-lactam antibiotics, the most widely used class in clinical practice. Antarctic soils harbor thousands of putative β-lactamase genes, yet the functional properties and resistance contributions of these enzymes remain largely unexplored. Here, we investigated the distribution, diversity, and functional impact of intrinsic class C β-lactamases (AmpC-type) in Antarctic members of the *Pseudomonas fluorescens* species complex, a lineage widely distributed in Antarctic soils and increasingly recognized as an opportunistic pathogen of humans, animals, and plants. Genome-scale analyses revealed that class C β-lactamases are intrinsic, widespread, and highly diverse within this lineage. Phenotypic assays demonstrated that Antarctic isolates exhibit elevated resistance to β-lactam antibiotics, particularly oxyimino-cephalosporins such as cefotaxime and ceftazidime, and that this phenotype is largely attenuated by β-lactamase inhibition. Heterologous expression of selected Antarctic AmpC variants in a susceptible *Escherichia coli* host confirmed their ability to increase minimum inhibitory concentrations to oxyimino-cephalosporins. Notably, several Antarctic β-lactamases harbor amino acid substitutions previously associated with extended-spectrum AmpC (ESAC) variants, including M174L and N346I, while others display broader substrate profiles despite lacking known ESAC-associated signatures. Comparative analyses further showed that the sequence diversity of Antarctic class C β-lactamases exceeds that reported for clinical PDC variants from *Pseudomonas aeruginosa*. Together, these findings provide functional evidence that pristine Antarctic environments constitute reservoirs of naturally occurring class C β-lactamases with extended-spectrum potential. Our results highlight the evolutionary depth and functional diversity of environmental β-lactamases and underscore the importance of incorporating remote ecosystems into One Health–oriented antimicrobial resistance surveillance frameworks.

## INTRODUCTION

Antimicrobial resistance (AMR) constitutes one of the most pressing global public health challenges of the 21st century. Antibiotic-resistant infections already account for a substantial burden of morbidity and mortality worldwide, and recent projections estimate that AMR-related deaths could surpass those caused by cancer by 2050 (Naghavi et al., 2024). Addressing this crisis requires integrative strategies that extend beyond clinical settings. In this context, the One Health framework has emerged as a central paradigm, emphasizing the interconnectedness of human, animal, and environmental health, and recognizing these ecosystems as reservoirs of antimicrobial determinants with potential clinical relevance (Larsson et al., 2023).

Among antimicrobial agents, β-lactam antibiotics remain the most widely prescribed class in clinical practice due to their broad spectrum of activity and favorable safety profiles (Mora-Ochomogo & Lohans, 2021). Consequently, resistance mechanisms targeting β-lactams are of particular concern. In Gram-negative bacteria, the production of β-lactamases is the predominant mechanism conferring resistance to these antibiotics (Bush, 2018). β-lactamases form a highly diverse group of enzymes that have evolved through ancient evolutionary processes as well as more recent selective pressures and are broadly distributed across both clinical and environmental bacterial populations (Bush & Bradford, 2020; Gholipour et al., 2024).

Within the genus *Pseudomonas*, resistance to β-lactam antibiotics is notably multifactorial, arising from the combined action of intrinsic β-lactamases, reduced outer membrane permeability, and multidrug efflux systems (Glen & Lamont, 2021). While *Pseudomonas aeruginosa* has been extensively studied as a major opportunistic pathogen, increasing attention has been directed toward non-*aeruginosa* species, particularly members of the *Pseudomonas fluorescens* species complex (Ishii et al., 2024; Salvà-Serra et al., 2025; Silverio et al., 2022; Thene et al., 2025). This complex comprises more than 50 validly described species characterized by remarkable ecological versatility, inhabiting soil, freshwater, marine environments, plants, animals, and, in some cases, human-associated niches (Garrido-Sanz et al., 2016). Importantly, several *P. fluorescens* complex species have been reported to display high levels of intrinsic resistance to β-lactams, even in environments with minimal or no direct anthropogenic antibiotic exposure (Silverio et al., 2022, 2025).

Antarctic ecosystems provide a particularly informative context in which to investigate intrinsic antimicrobial resistance. Despite their relative isolation and low human impact, Antarctic soils harbor bacterial communities enriched in antibiotic resistance genes, including thousands of putative β-lactamase sequences (Van Goethem et al., 2018; Marcoleta et al., 2022; Coche-Miranda et al., 2025). Recent studies have reported Antarctic *P. fluorescens* complex isolates exhibiting elevated resistance to multiple β-lactam antibiotics, in some cases surpassing resistance levels observed in the reference strain *P. aeruginosa* PAO1 (García-Laviña et al., 2023; Marcoleta et al., 2022; Silverio et al., 2025). These observations suggest that β-lactam resistance in Antarctic *Pseudomonas* is intrinsic rather than acquired through recent horizontal gene transfer from clinical sources (Van Goethem et al., 2018). Genomic analyses have implicated chromosomally encoded class C β-lactamases (AmpC-type) and multidrug efflux systems, such as MexAB-OprM, as potential contributors to these resistance phenotypes (García-Laviña et al., 2023).

Class C β-lactamases (AmpC) are clinically relevant enzymes that confer resistance to a range of β-lactam antibiotics, including penicillins and several cephalosporins (Jacoby, 2009). In *P. aeruginosa*, AmpC-mediated resistance commonly arises through *ampC* overexpression or structural modifications that enhance catalytic efficiency against antipseudomonal β-lactams (Glen & Lamont, 2021). In particular, *Pseudomonas*-derived cephalosporinase (PDC) variants with extended-spectrum activity toward oxyimino-cephalosporins, such as ceftazidime, cefotaxime, and cefepime, have been increasingly reported in clinical isolates, likely as a consequence of mutations driven by antibiotic selective pressure (Berrazeg et al., 2015; Colque et al., 2022; Rodríguez-Martínez et al., 2009). These extended-spectrum AmpC variants represent a significant therapeutic challenge, as they compromise the efficacy of key antipseudomonal agents.

Despite the growing recognition of extended-spectrum class C β-lactamases in clinical *P. aeruginosa*, the diversity, functional properties, and contribution to resistance of class C β-lactamases from environmental *Pseudomonas* species, particularly non-*aeruginosa* lineages, remain poorly characterized. This gap is especially relevant for members of the *P. fluorescens* species complex, which are increasingly detected in both environmental and clinical contexts. In particular, it remains unclear whether the sequence diversity of AmpC beta-lactamases encoded by environmental *Pseudomonas* represents neutral genomic variation or translates into different resistance profiles against clinically relevant beta-lactam antibiotics.

In this study, we examined class C beta-lactamases encoded by Antarctic *P. fluorescens* complex bacteria using an integrated approach combining comparative genomics with phenotypic and molecular analyses. Specifically, we sought to clarify the evolutionary distribution of these enzymes within the lineage and to assess their potential role in shaping resistance to clinically relevant β-lactam antibiotics. By focusing on bacteria originating from pristine Antarctic environments, this work provides insight into the environmental reservoirs and functional potential of class C beta-lactamases and highlights the relevance of environmental lineages for One Health-oriented perspectives on antimicrobial resistance.

## RESULTS

### Class C β-lactamases are intrinsic, diverse, and widespread in Antarctic *Pseudomonas fluorescens* complex bacteria

To determine whether class C β-lactamases represent sporadic traits or intrinsic features of Antarctic *P. fluorescens* complex bacteria, we first examined their distribution at the genomic scale. A total of 2,346 *Pseudomonas* genomes were analyzed, including 120 genomes derived from Antarctic isolates. Based on average nucleotide identity (ANI) values and phylogenetic placement relative to representative species of the *P. fluorescens* complex, 737 genomes were classified as members of this lineage, of which 104 corresponded to Antarctic *Pseudomonas* spp. (Figure S1).

Within this Antarctic subset, 100 class C β-lactamase (*ampC*-type) genes were identified, indicating that these enzymes are nearly ubiquitous and therefore intrinsic to Antarctic *P. fluorescens* complex bacteria (Table S1). This widespread distribution supports the idea that class C β-lactamases constitute a core genomic feature of this lineage rather than resistance determinants recently acquired through horizontal gene transfer.

The 100 Antarctic AmpC-type enzymes exhibited a broad range of amino acid sequence identity, spanning from 49% to 98%, which is substantially greater than the diversity reported for *Pseudomonas*-derived cephalosporinase (PDC) variants from the *P. aeruginosa* lineage, which typically share 87-100% identity. When stratified by phylogenetic subgroup, sequences from the *P. mandelii* subgroup displayed the greatest diversity (57.8-96.6%), followed by those from the *P. fluorescens* and *P. fragi* subgroups, indicating pronounced diversification within Antarctic *P. fluorescens* complex bacteria.

To further examine evolutionary relationships among the Antarctic class C β-lactamase sequences, they were clustered at 90% amino acid identity, yielding 35 representative sequences that were analyzed together with the PDC-1 reference enzyme from *P. aeruginosa* PAO1 (Table S2). Phylogenetic analysis revealed that Antarctic class C β-lactamases segregate into four major clades corresponding to the four recognized subgroups of the *P. fluorescens* species complex (Figure 1). Sequences belonging to the *P. mandelii* and *P. fluorescens* subgroups were each divided into two major clades, reflecting substantial intra-subgroup diversification, whereas sequences from the *P. fragi* subgroup formed a distinct and highly divergent monophyletic clade.

**Figure 1.**
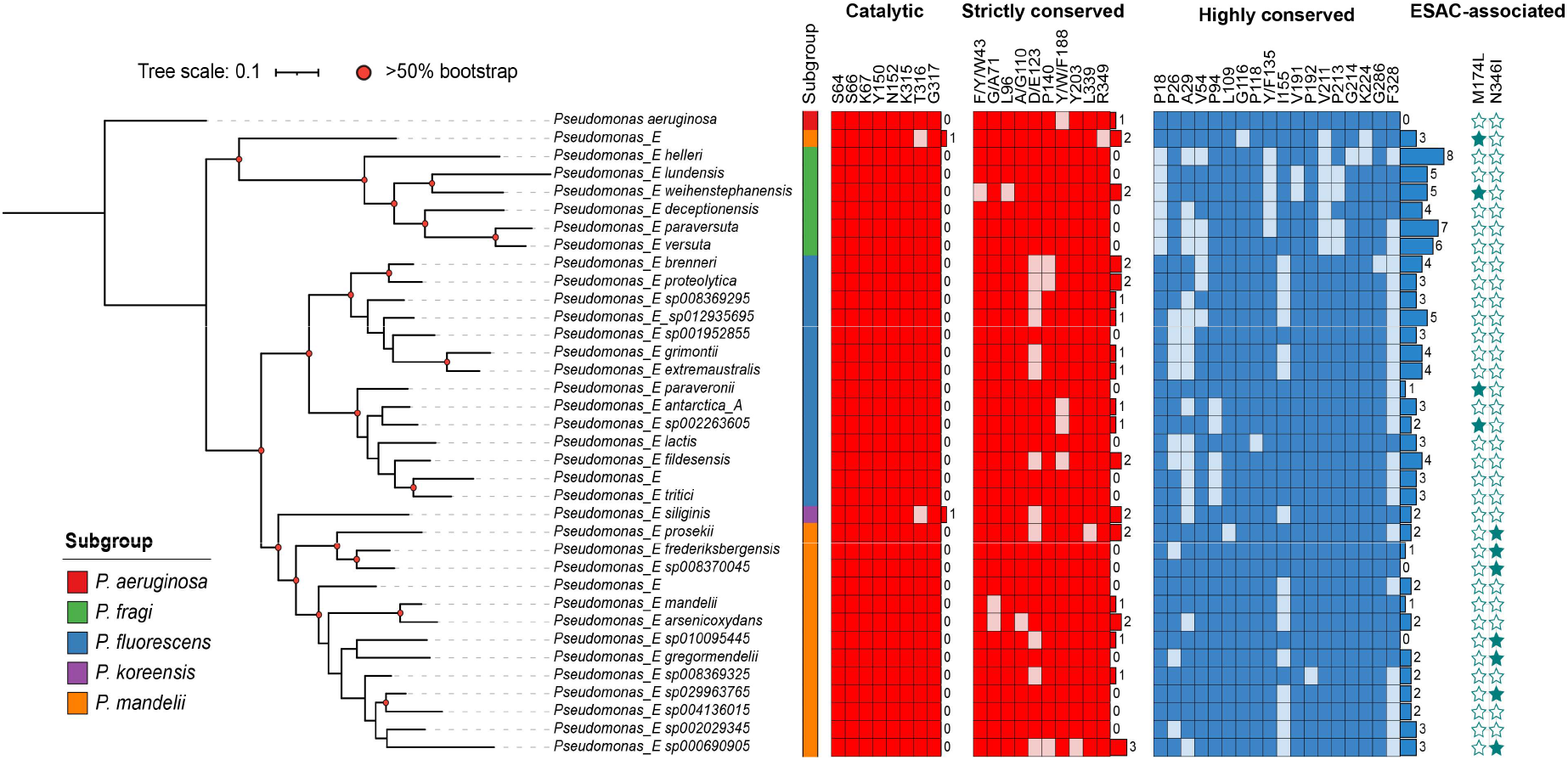
Genome-scale distribution, diversity, and phylogenetic relationships among class C β-lactamases in Antarctic *Pseudomonas fluorescens* complex bacteria. Phylogenetic analysis of representative class C β-lactamase (AmpC-type) sequences identified in Antarctic Pseudomonas fluorescens complex genomes, analyzed together with the PDC-1 reference enzyme from Pseudomonas aeruginosa PAO1. Sequence conservation and molecular features are summarized using the structural alignment-based numbering (SANC) system, including catalytic residues, strictly conserved residues, highly conserved residues, and the presence of extended-spectrum AmpC (ESAC)-associated substitutions (M174L and N346I).

Despite this pronounced sequence diversity, Antarctic class C β-lactamases largely retained residues required for catalytic activity. Conservation analysis based on the structural alignment-based numbering (SANC) system showed that 34 of the 36 representative sequences conserved 7 of the 8 catalytic residues (Table S3). The remaining two sequences carried substitutions at position T316 (T316S or T316P), a position previously reported to tolerate variation without loss of β-lactamase function. Analysis of strictly and highly conserved residues further indicated high overall conservation across the dataset.

Finally, we assessed the occurrence of amino acid substitutions previously associated with extended-spectrum AmpC (ESAC) variants. Among the Antarctic class C β-lactamases, several sequences carried substitutions corresponding to M174L or N346I, which have been reported in extended-spectrum AmpC variants from clinical *Pseudomonas* isolates. These substitutions were detected in multiple Antarctic genomes originating from pristine environments, indicating that ESAC-associated sequence features are present at the population level within Antarctic *P. fluorescens* complex bacteria (Figure 1).

Together, these genome-scale analyses demonstrate that Antarctic *P. fluorescens* complex bacteria harbor an intrinsically encoded and highly diverse repertoire of class C β-lactamases. This diversity provides a genomic framework for exploring whether these enzymes contribute to β-lactam resistance phenotypes in native Antarctic isolates.

### Antarctic *Pseudomonas fluorescens* complex strains exhibit elevated intrinsic resistance to β-lactam antibiotics

Antarctic *P. fluorescens* complex strains have been reported to display elevated intrinsic resistance to β-lactam antibiotics, even in environments with minimal anthropogenic antibiotic exposure. Given the widespread and diverse repertoire of class C β-lactamases identified at the genomic level in Antarctic *P. fluorescens* complex bacteria, we asked whether this genetic potential translates into measurable resistance phenotypes in native Antarctic isolates.

To address this question, eight Antarctic strains belonging to six species within the complex were selected for phenotypic characterization, spanning multiple phylogenetic subgroups (Table S4). β-lactam susceptibility profiles were first evaluated by disk diffusion using a panel of 16 β-lactam antibiotics. Several Antarctic strains displayed higher levels of resistance than the reference strain *P. aeruginosa* PAO1, which harbors well-characterized intrinsic β-lactam resistance mechanisms typical of this species (Grace et al., 2022) (Figure 2). Notably, four of the eight Antarctic strains exhibited reduced susceptibility across multiple β-lactams relative to PAO1. Resistance to clinically relevant antipseudomonal β-lactams was frequently observed. Seven of the eight Antarctic strains were resistant to the monobactam aztreonam, two were resistant to the extended-spectrum cephalosporin ceftazidime, and two were resistant to the carbapenem meropenem.

**Figure 2.**
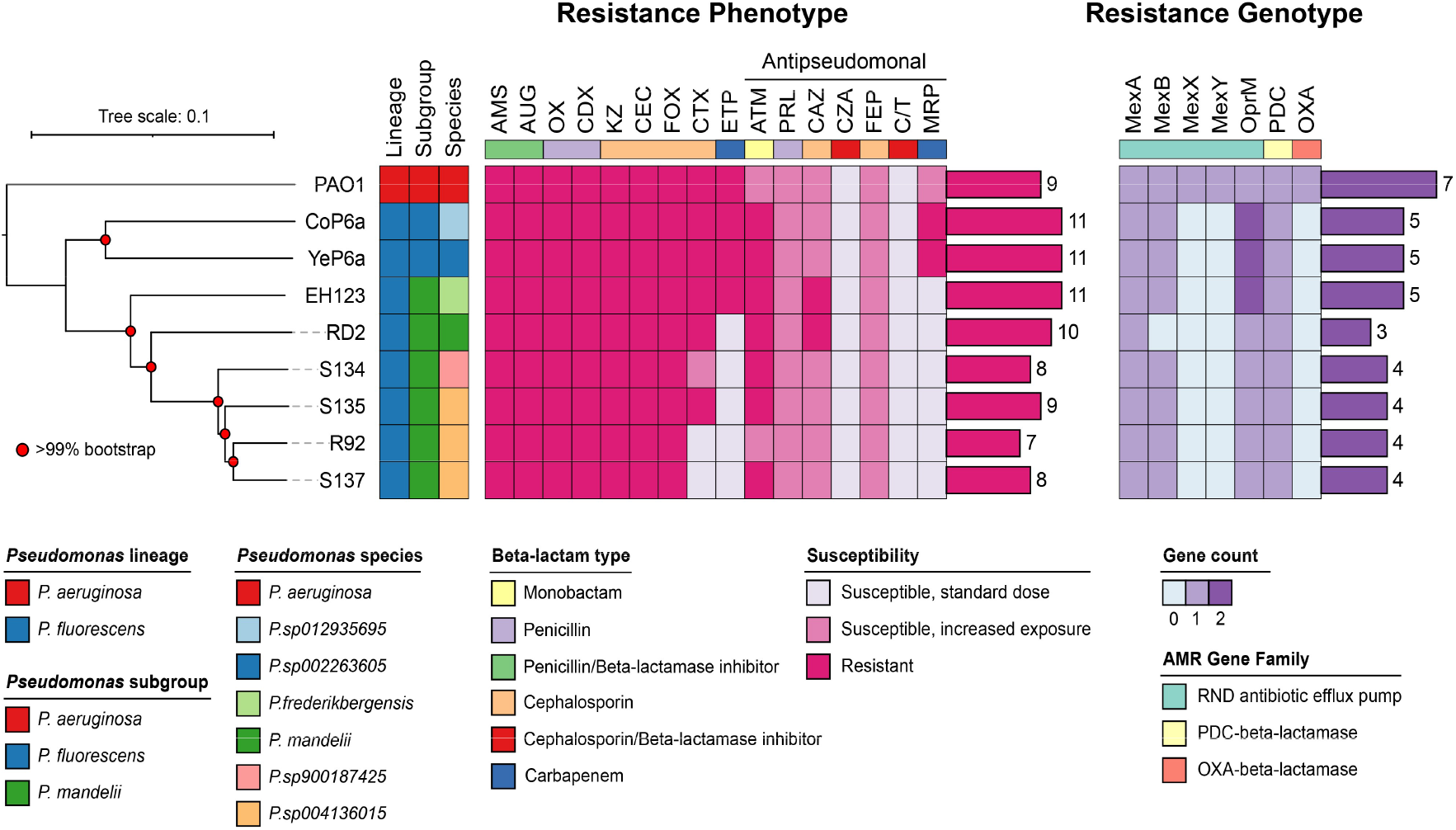
Phenotypic and genotypic profiles of β-lactam resistance in Antarctic *Pseudomonas fluorescens* complex strains and *Pseudomonas aeruginosa* PAO1. β-lactam susceptibility was assessed by disk diffusion assays in eight Antarctic *P. fluorescens* complex strains and the reference strain *P. aeruginosa* PAO1. Predicted β-lactam resistance determinants were identified using the RGI-CARD platform. The figure presents susceptibility patterns together with the distribution of major β-lactam resistance gene families, including class C β-lactamases and efflux systems.

To assess the contribution of β-lactamase activity to these resistance phenotypes, ceftazidime susceptibility was also evaluated in the presence of the β-lactamase inhibitor avibactam. In all Antarctic strains, avibactam increased susceptibility to ceftazidime, indicating that β-lactamase activity is a major mechanism contributing to resistance to this antibiotic (Figure 2).

To place these phenotypic observations in a genomic context, β-lactam resistance determinants were predicted using the RGI-CARD platform. Overall, Antarctic *P. fluorescens* complex strains harbored fewer predicted β-lactam resistance genes than *P. aeruginosa* PAO1 (Figure 2). In particular, they lacked orthologs of the MexXY efflux pump system and the OXA-like β-lactamase poxB. In contrast, all Antarctic strains encoded a chromosomal PDC-like class C β-lactamase, and seven of the eight strains encoded the MexAB-OprM efflux system. Notably, strains CoP6a, YeP6b, and EH123 carried two copies of the *oprM* gene. Together, these observations suggest that elevated β-lactam resistance in Antarctic *P. fluorescens* strains is largely mediated by intrinsic resistance mechanisms, with class C β-lactamases representing a central component.

To further characterize this resistance phenotype quantitatively, broth microdilution assays were performed to determine minimum inhibitory concentrations (MICs) for five β-lactam antibiotics in the presence or absence of avibactam (Table 1). The four most resistant Antarctic strains identified by disk diffusion, CoP6a, YeP6b, EH123, and RD2, were selected for this analysis. MIC values for ampicillin were similarly high in all Antarctic strains compared with *P. aeruginosa* PAO1. In contrast, MICs for cefotaxime and ceftazidime were 8 to 16-fold and 4 to 16-fold higher than those of PAO1, respectively, with strains EH123 and RD2 exhibiting the highest MIC values. For cefepime and meropenem, differences were more modest, with MIC values ranging from equal to those of PAO1 to up to fourfold higher. In this respect, YeP6b and CoP6a showed the highest MICs for meropenem.

**Table 1.** Minimum inhibitory concentrations (MICs) of β-lactam antibiotics for *Pseudomonas aeruginosa* PAO1 and Antarctic *Pseudomonas fluorescens* complex strains, determined in the presence or absence of avibactam.

| Strain | MIC (ug/mL) |  |  |  |  |  |  |  |  |  |
| --- | --- | --- | --- | --- | --- | --- | --- | --- | --- | --- |
|  | AMP | AMP-AVI | CTX | CTX-AVI | CAZ | CAZ-AVI | FEP | FEP-AVI | MRP | MRP-AVI |
| PAO1 | 256 | 64 | 8 | 8 | 1 | 1 | 0.5 | 0.5 | 1 | 1 |
| CoP6a | >256 | 256 | >256 | 64 | 8 | 8 | 8 | 2 | 8 | 2 |
| EH123 | >256 | 256 | >256 | <8 | 32 | <2 | 2 | 0.5 | <0.25 | 0.25 |
| RD2 | >256 | 128 | 128 | <8 | 16 | <2 | 8 | 1 | 2 | 1 |
| YeP6b | >256 | 256 | 128 | 32 | 8 | 4 | 4 | 1 | 4 | 1 |

In the presence of avibactam, MIC values for ampicillin remained unchanged in all strains, whereas MICs for cefotaxime decreased by at least fourfold in all cases. For ceftazidime, MIC values decreased by at least twofold in all strains except CoP6a, while for meropenem, a similar reduction was observed in all strains except EH123. These results indicate that PDC-like class C β-lactamases primarily mediate resistance to the oxyimino-cephalosporins cefotaxime and ceftazidime in Antarctic *P. fluorescens* complex strains, with a more limited contribution to resistance against cefepime and meropenem. This effect was most pronounced in strains exhibiting the highest MIC values, such as EH123 and RD2.

Together, these findings demonstrate that Antarctic *P. fluorescens* complex strains exhibit elevated intrinsic resistance to β-lactam antibiotics and identify class C β-lactamases as a major contributor to cephalosporin resistance in their native genomic context. These observations provide the rationale for examining the molecular features of selected Antarctic class C β-lactamases in greater detail.

### Molecular features and extended-spectrum associated signatures in Antarctic class C β-lactamases

Given that Antarctic *P. fluorescens* complex strains exhibit elevated resistance to oxyimino-cephalosporins that is largely attenuated by β-lactamase inhibition, we next examined the molecular characteristics of their class C β-lactamases. Specifically, we focused on the enzymes selected for functional characterization to assess sequence conservation, diversity, and the presence of amino acid substitutions previously associated with extended-spectrum AmpC activity.

The four Antarctic class C β-lactamases analyzed here, derived from strains EH123, RD2, CoP6a, and YeP6b, shared 65.8 to 73.5 percent amino acid sequence identity among themselves and 61.3 to 62.6 percent identity with the reference PDC-1 β-lactamase from *P. aeruginosa* PAO1 (Figure 3A; Table S5). Comparison with the BLDB database further showed that these enzymes shared 62.95 to 83.59 percent amino acid sequence identity with their closest characterized homologs, corresponding to the PFL-4, PFL-5, and PFL-6 variants previously associated with *P. fluorescens* strains.

**Figure 3.**
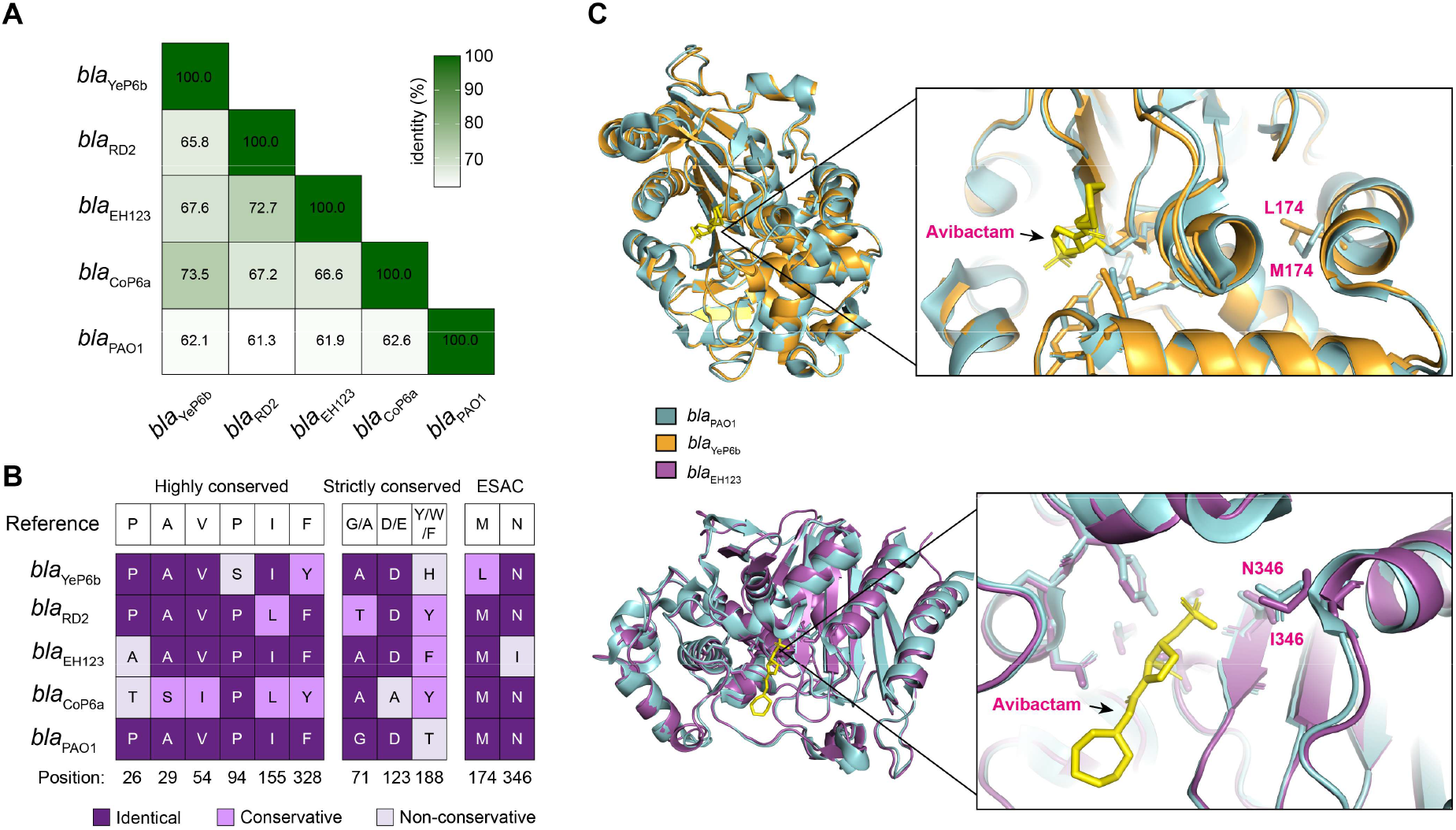
Molecular and structural features of Antarctic class C β-lactamases. (A) Amino acid sequence identity matrix comparing Antarctic β-lactamases with the PDC-1 reference sequence from *Pseudomonas aeruginosa* PAO1. (B) Conservation analysis of selected residues across five analyzed β-lactamases. The first two panels indicate the conservation status of six highly conserved residues and three strictly conserved residues. The final panel shows the presence or absence of the M174L and N346I substitutions previously associated with extended-spectrum AmpC (ESAC) activity. (C) Structural localization of ESAC-associated substitutions. Upper panel: position of the M174L substitution in the AlphaFold-predicted structure of blaYeP6b, superimposed on the PDC-1 reference structure in complex with the β-lactamase inhibitor avibactam (PDB: 2WZZ). Lower panel: position of the N346I substitution in the AlphaFold-predicted structure of blaEH123, superimposed on the same PDC-1 reference structure.

Following the observed sequence divergence among Antarctic class C β-lactamases, we next examined whether this variability was accompanied by conservation of molecular features required for enzymatic activity. All four Antarctic β-lactamases conserved the canonical class C active-site motifs S64XSK, Y150XN, and K315TG (Figure S2), indicating preservation of the core catalytic architecture despite substantial sequence divergence.

To further assess molecular conservation, residues conserved among class C β-lactamases were analyzed following the classification proposed by Philippon et al. (2022), which distinguishes 48 strictly conserved residues and 34 highly conserved residues. Across the four Antarctic enzymes, 98 percent of strictly conserved residues and between 85 and 100 percent of highly conserved residues were retained (Table S6). These results indicate that, although Antarctic class C β-lactamases are divergent at the primary sequence level, they largely maintain the conserved residue framework characteristic of functional AmpC enzymes.

Within this overall pattern of conservation, differences among the four enzymes were not uniformly distributed. Among them, blaCoP6a exhibited the highest number of substitutions within the highly conserved residue category (Table S6). Specifically, the Antarctic β-lactamases differed at six positions classified as highly conserved, namely P26, A29, V54, P94, I155, and F328, as well as at three positions classified as strictly conserved, G71A, D123E, and Y188W or F. While most substitutions were biochemically conservative, several nonconservative changes were observed, including P94S in blaYeP6b, P26A in blaEH123, and P26T in blaCoP6a (Figure 3B). These substitutions highlight discrete molecular differences among Antarctic enzymes that may contribute to functional variability.

Given the elevated resistance to oxyimino-cephalosporins observed in native Antarctic strains, we next examined whether the Antarctic β-lactamases carried amino acid substitutions previously associated with extended-spectrum AmpC variants. In this analysis, blaYeP6b was found to harbor the M174L substitution, whereas blaEH123 carried the N346I substitution (Figure 3B and 3C), both of which have been implicated in extended-spectrum activity in class C β-lactamases from Pseudomonas aeruginosa. The M174L substitution is located within the H6 helix, outside the active site, whereas N346I lies within the H11 helix, in proximity to the active-site region (Figure 3C). The presence of these substitutions in Antarctic enzymes provides a molecular link between the genomic diversity described in Section 1 and the resistance phenotypes observed in Section 2.

Together, these analyses indicate that Antarctic class C β-lactamases combine substantial sequence divergence with strong conservation of residues essential for catalytic activity and structural integrity, while also harboring specific substitutions previously associated with extended-spectrum AmpC variants. These molecular features provide a clear rationale for experimentally testing whether Antarctic class C β-lactamases can intrinsically mediate resistance to extended-spectrum β-lactam antibiotics when expressed in a heterologous host.

### Antarctic class C β-lactamases display extended-spectrum activity in a heterologous host

Building on the molecular features identified in Antarctic class C β-lactamases and their association with elevated resistance phenotypes in native strains, we next examined whether these enzymes can mediate resistance to β-lactam antibiotics independent of their native genomic background. To this end, selected Antarctic class C β-lactamase genes were heterologously expressed in a β-lactam-susceptible *E. coli* host, together with the PDC-1 reference enzyme from *P. aeruginosa* PAO1, and their contribution to β-lactam susceptibility was evaluated.

Recombinant *E. coli* strains expressing Antarctic class C β-lactamases from strains EH123, RD2, CoP6a, and YeP6b, as well as the PDC-1 reference enzyme from *P. aeruginosa* PAO1, were first evaluated by disk diffusion assays against a panel of β-lactam antibiotics. Reduced susceptibility relative to the control strain carrying the empty vector was used as an indicator of β-lactamase activity, enabling a comparative assessment of substrate profiles across the different enzymes.

Using this criterion, all recombinant strains expressing Antarctic class C β-lactamases, as well as the strain expressing the PDC-1 reference enzyme, exhibited β-lactamase activity against at least 7 of the 18 β-lactam antibiotics tested (Figure 4). The observed activity was primarily directed against penicillins and first- and second-generation cephalosporins, consistent with the expected substrate profile of class C β-lactamases (Philippon et al., 2022). Notably, the β-lactamase derived from the Antarctic strain RD2 displayed a broader overall activity spectrum than the PDC-1 control enzyme.

**Figure 4.**
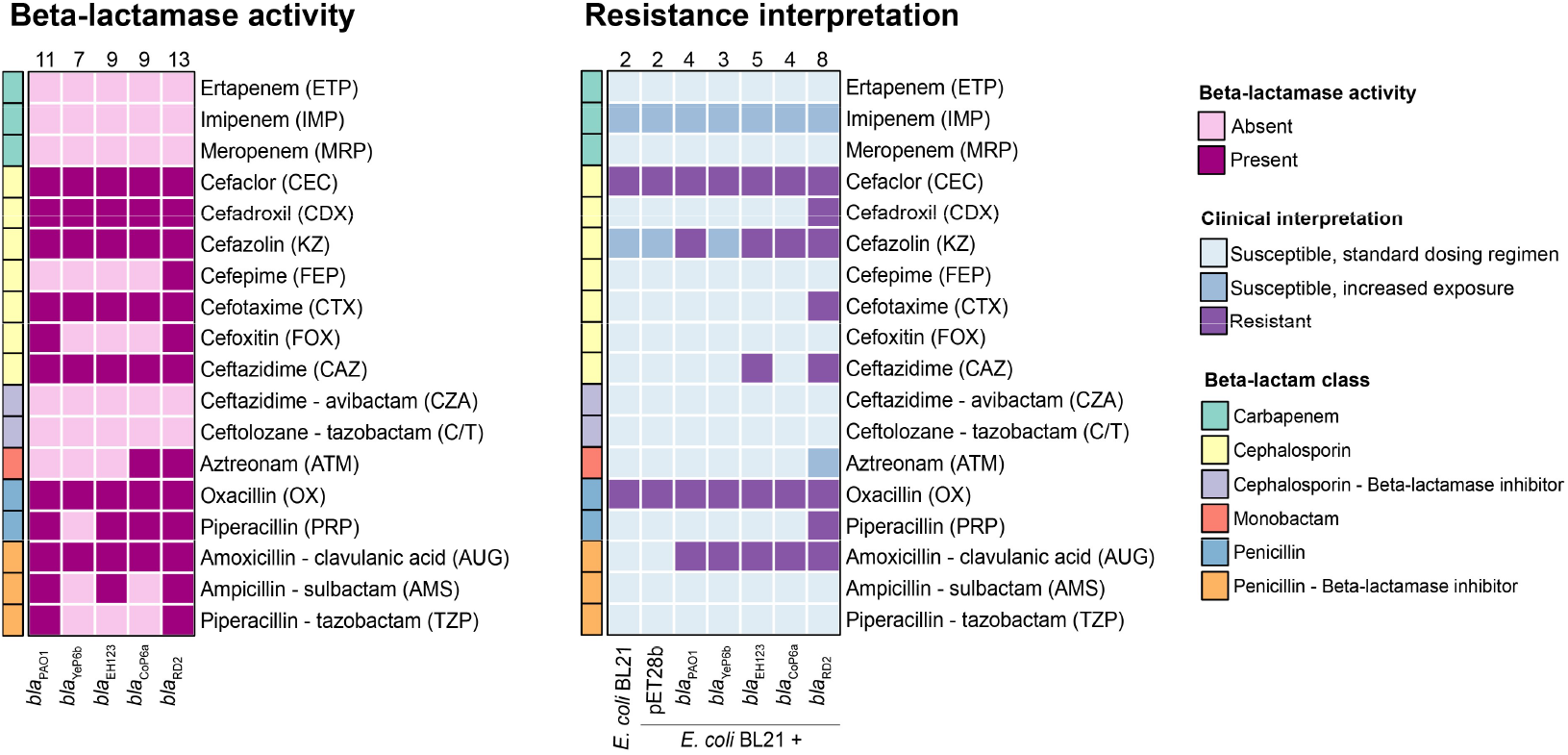
Antibiotic susceptibility profiles of *E. coli* BL21 strains expressing Antarctic class C β-lactamases. (left panel) Matrix summarizing β-lactamase activity based on disk diffusion assays. Activity was defined as a reduction in inhibition zone diameter of at least 5 mm relative to the control strain *E. coli* BL21 carrying the empty pET28b vector. (right panel) Interpretation of inhibition zone diameters according to EUCAST clinical breakpoints for Enterobacterales.

When disk diffusion results were interpreted using EUCAST clinical breakpoints for Enterobacterales, all Antarctic β-lactamases and the PDC-1 reference enzyme conferred resistance to amoxicillin-clavulanic acid, followed by cefazolin, except for the blaYeP6b β-lactamase. Cefaclor and oxacillin were excluded from this analysis because *E. coli* is intrinsically resistant to these antibiotics. Importantly, the Antarctic blaEH123 and blaRD2 β-lactamases conferred resistance to the extended-spectrum oxyimino-cephalosporin ceftazidime. In addition, blaRD2 conferred resistance to cefadroxil, cefotaxime, and piperacillin, as well as intermediate susceptibility to aztreonam. Notably, piperacillin and aztreonam are clinically relevant antipseudomonal β-lactam antibiotics.

To further quantify these susceptibility patterns, broth microdilution assays were performed to determine minimum inhibitory concentrations for five selected β-lactam antibiotics (Table 2). Expression of Antarctic class C β-lactamases resulted in reduced susceptibility to ampicillin, cefotaxime, and ceftazidime, whereas susceptibility to cefepime and meropenem was largely unchanged. Notably, all recombinant strains expressing Antarctic β-lactamases exhibited higher MIC values for ceftazidime than the strain expressing the PDC-1 reference enzyme from *P. aeruginosa* PAO1.

**Table 2.** Minimum inhibitory concentrations of β-lactam antibiotics for recombinant *Escherichia coli* BL21 strains expressing Antarctic class C β-lactamases and the PDC-1 reference enzyme from *Pseudomonas aeruginosa* PAO1.

| Strain | MIC (μg/mL) | | | | |
| --- | --- | --- | --- | --- | --- |
|  | AMP | CTX | CAZ | FEP | MRP |
| *E. coli* BL21 | <2 | <0.125 | <0.125 | <0.125 | 0.025 |
| *E. coli* BL21-pET28b | <2 | <0.125 | <0.125 | <0.125 | 0.025 |
| blaPAO1 | 64 | 4 | 0.5 | <0.125 | 0.025 |
| blaYeP6b | 16 | 0.5 | 2 | <0.125 | 0.025 |
| blaEH123 | 32 | 4 | 4 | <0.125 | 0.025 |
| blaCoP6a | 16 | 4 | 1 | <0.125 | 0.025 |
| blaRD2 | 64 | >4 | 4 | <0.125 | 0.025 |

To confirm that the observed resistance phenotypes were mediated by class C β-lactamase activity, ampicillin, cefotaxime, and ceftazidime were tested in combination with the β-lactamase inhibitor avibactam against selected recombinant strains, including those expressing blaEH123 and blaRD2, together with the blaPAO1 reference enzyme (Table S7). In all cases, MIC values were restored in the presence of avibactam, supporting that these Antarctic class C β-lactamases are effectively inhibited by avibactam.

Together, these results demonstrate that Antarctic class C β-lactamases can intrinsically mediate resistance to β-lactam antibiotics when expressed in a susceptible heterologous host, including resistance to extended-spectrum oxyimino-cephalosporins such as cefotaxime and ceftazidime. This functional activity is consistent with the molecular features identified in these enzymes and confirms that Antarctic environments harbor naturally occurring class C β-lactamases with extended-spectrum potential.

## DISCUSSION

Understanding how antimicrobial resistance determinants are distributed and function in natural environments is essential for clarifying their evolutionary origins and potential clinical implications. Multiple studies have documented elevated tolerance to β-lactam antibiotics among environmental *Pseudomonas* strains, including members of the *P. fluorescens* species complex isolated from Antarctic ecosystems. However, the specific contribution of class C β-lactamases to these patterns, and whether naturally occurring variants exhibit extended-spectrum properties, has remained unresolved. Here, we addressed these questions by integrating genome-scale analyses with phenotypic and molecular characterization of Antarctic *P. fluorescens* complex isolates.

Early insight into Antarctic class C β-lactamases came from the characterization of PFL-P1, the first enzyme described from an Antarctic *P. fluorescens* strain. PFL-P1 was reported to possess fewer hydrogen bonds, a lower proline content, and a reduced proportion of arginine relative to lysine residues, structural features proposed to confer increased flexibility and potentially explain its activity toward bulky penicillins (Michaux et al., 2008; Pierrard, 1998). However, beyond these structural and kinetic observations, its contribution to the antibiotic susceptibility profile of the native host, as well as its activity against clinically relevant antipseudomonal β-lactams, remained unclear. Thus, although individual Antarctic enzymes had been described, their functional impact at the strain level had not been systematically examined.

Subsequent reports documented elevated β-lactam tolerance among Antarctic *Pseudomonas* isolates, reinforcing the need to clarify the mechanisms involved. García-Laviña et al. (2023) reported that *Pseudomonas* sp. AU10 was resistant to aminopenicillins, cefotaxime, and aztreonam and carried both an AmpC-type β-lactamase and the MexAB efflux pump, although their relative contributions were not determined. Similarly, Silverio et al. (2025) identified *P. fluorescens* complex strains displaying resistance or intermediate susceptibility to aztreonam, ceftazidime, cefepime, and, less frequently, imipenem. In the present study, Antarctic isolates also exhibited reduced susceptibility to antipseudomonal β-lactams such as aztreonam and ceftazidime, although none were resistant to cefepime or meropenem. Differences across studies may be attributable to ecological context and sampling source. The isolates analyzed here were largely obtained from minimally impacted environments, whereas those examined by Silverio et al. (2025) included strains associated with plants or animals, which may experience distinct host-related or competitive selective pressures. Importantly, our results indicate that class C β-lactamases play a central role in resistance to cefotaxime and ceftazidime in the strains examined here, underscoring the need to evaluate their contribution across a broader range of Antarctic isolates. To further explore the mechanistic basis of this effect, we compared the resistance profiles of Antarctic strains with those of the well-characterized reference strain *P. aeruginosa* PAO1.

Clear differences emerged between Antarctic isolates and PAO1. In Antarctic strains, class C β-lactamases were primarily associated with resistance to cefotaxime and ceftazidime, whereas inhibition of β-lactamase activity had minimal impact on ampicillin susceptibility. PAO1, by contrast, displayed a distinct response pattern. These differences cannot be explained solely by catalytic properties, as all enzymes conferred comparable reductions in ampicillin susceptibility when expressed in *E. coli*, yet differed in their effects on ceftazidime. The influence of AmpC on antibiotic response reflects both enzymatic activity and regulatory control. Penicillins are generally strong inducers and efficient substrates of AmpC enzymes (Jacoby, 2009), a property partially supported by heterologous expression assays. However, *ampC* inducibility was not directly measured in the Antarctic strains analyzed here. Nonetheless, variation in regulatory response may contribute to the distinct profiles observed between Antarctic isolates and PAO1. In support of this possibility, the Antarctic strain *Pseudomonas* sp. IB20, belonging to the same species as YeP6b, was recently shown to exhibit strong *ampC* induction in response to ampicillin, with an approximately 510-fold increase in expression, whereas induction in PAO1 was comparatively modest (Higuera-Llantén et al., 2024). These findings suggest that differences in *ampC* regulation may influence β-lactam susceptibility in environmental strains.

Despite this, inhibition of class C β-lactamase activity with avibactam did not alter ampicillin susceptibility in the Antarctic isolates examined here. This indicates that, for penicillins, AmpC activity alone is unlikely to account for the observed resistance levels. Instead, additional intrinsic mechanisms, including the MexAB-OprM efflux system identified in most strains, likely play a dominant role in mediating resistance to ampicillin.

This substrate-dependent pattern became more apparent when examining oxyimino-cephalosporins. In contrast to the limited impact observed for penicillins, inhibition of β-lactamase activity resulted in a consistent reduction in MIC values for cefotaxime and ceftazidime in Antarctic strains, whereas no comparable effect was observed in PAO1. Although ceftazidime is considered a weak inducer of *ampC* expression, it can be hydrolyzed when sufficient enzyme is present (Jacoby, 2009). While induction levels were not quantified in this study, expression of Antarctic AmpC enzymes in *E. coli* was associated with higher ceftazidime MIC values than those observed for PDC-1, suggesting differences in substrate profile or catalytic performance. These intrinsic properties likely contribute to the enhanced cephalosporin resistance observed in native Antarctic strains.

Although the effect of AmpC was most pronounced for cefotaxime and ceftazidime, class C β-lactamases also appeared to contribute to reduced susceptibility to cefepime and meropenem in some Antarctic strains. Notably, this effect was not recapitulated in recombinant *E. coli* expressing the same enzymes. This discrepancy suggests that the intrinsic catalytic activity of Antarctic AmpC variants against cefepime and meropenem may be modest but becomes phenotypically relevant in the native host background. In Antarctic strains, the combined influence of β-lactamase activity, regulatory dynamics, outer membrane permeability, and efflux systems may collectively elevate resistance levels. These observations underscore the importance of interpreting β-lactam susceptibility in *Pseudomonas* as the outcome of interacting intrinsic mechanisms rather than the effect of a single determinant (Glen & Lamont, 2021; Livermore, 1995). Given the measurable differences in substrate profiles among Antarctic AmpC variants, it is therefore pertinent to consider whether structural variation within these enzymes may confer extended-spectrum properties comparable to those described in clinical settings.

Several extended-spectrum PDC variants from *P. aeruginosa* have been associated with substitutions located in structurally relevant regions, including the omega loop, the R2 domain, and the C-terminal region of the enzyme (Berrazeg et al., 2015; Colque et al., 2022; Rodríguez-Martínez et al., 2009). In this study, we identified naturally occurring class C β-lactamase variants within the genus *Pseudomonas* that harbor substitutions previously linked to extended-spectrum AmpC enzymes.

Notably, *bla*_EH123_ carries the N346I substitution, which has been associated with extended-spectrum activity in clinical PDC variants (Berrazeg et al., 2015). This substitution has also been shown to enhance the activity of plasmid-borne CMY-2 and *Enterobacter* AmpC enzymes against ceftazidime and cefepime (Dahyot et al., 2013; Livermore et al., 2012). In our dataset, N346I was identified in seven Antarctic *P. fluorescens* complex sequences, representing approximately 20 percent of analyzed variants, suggesting that Antarctic environments may constitute a natural reservoir of ESAC-like enzymes. Similarly, *bla*_YeP6b_ carries the M174L substitution, previously associated with extended-spectrum PDC variants (Berrazeg et al., 2015). However, in the clinical variant PDC-82, this substitution occurs in conjunction with F121L, which is absent in *bla*_YeP6b_. Therefore, the extent to which M174L alone contributes to extended-spectrum activity remains unclear and warrants further functional validation.

Interpretation of ESAC-associated residues across highly divergent sequence backgrounds must be approached cautiously. Antarctic class C β-lactamases share limited sequence identity with clinical PDC enzymes, and the structural consequences of individual substitutions may vary depending on overall protein architecture. In this context, *bla*_RD2_ exhibited a broader substrate spectrum despite lacking previously described ESAC-associated residues, indicating that alternative structural determinants may underlie its extended activity profile. Detailed biochemical and structural studies will be required to resolve the mechanistic basis of these phenotypes.

Beyond their contribution to antibiotic susceptibility, the ecological and physiological roles of class C β-lactamases in Antarctic soils merit consideration. Bacteria capable of hydrolyzing β-lactam compounds are widely reported in these environments (Coche-Miranda et al., 2025), yet the selective forces maintaining this activity in remote ecosystems remain incompletely understood.

Serine β-lactamases have been proposed to contribute to cell-wall biosynthesis and remodeling, suggesting that their ancestral functions may extend beyond antibiotic inactivation (Jacobs et al., 1994; Massova & Mobashery, 1998). In this context, class C β-lactamases appear to have evolved relatively recently within the penicillin-binding protein (PBP)-like superfamily, potentially originating from AmpH-type low-molecular-weight PBPs or related esterases (Fröhlich et al., 2021). Indeed, the classification of certain carboxylesterases as bona fide β-lactamases remains under active discussion (Philippon et al., 2022), underscoring the blurred functional boundaries within this protein family. These evolutionary considerations indicate that catalytic activity toward β-lactam rings arose from physiological roles that predate modern antibiotic exposure.

Alternatively, β-lactamases may provide defense against naturally occurring antibiotic-producing microbes in soil ecosystems (Gholipour et al., 2025). β-lactam production by fungi has been documented in Antarctic soils (Pantůček et al., 2018), indicating that environmental bacteria may encounter selective pressures favoring β-lactam hydrolytic activity even in remote habitats. Furthermore, members of the *P. fluorescens* complex can colonize diverse animals and plants (Silverio et al., 2022), suggesting that these enzymes may enhance persistence in host-associated or polymicrobial environments. This ecological role has been demonstrated in *Pectobacterium versatile*, where β-lactamase production maintains the diversity of β-lactam-susceptible strains during potato infection (Lorang et al., 2025), illustrating how such enzymes can influence microbial community structure rather than merely confer individual resistance.

In summary, this study demonstrates that Antarctic *P. fluorescens* complex bacteria encode intrinsic and highly diverse class C β-lactamases that contribute substantially to reduced susceptibility against clinically relevant β-lactam antibiotics and include naturally occurring variants with extended-spectrum potential. These findings support the view that pristine and remote environments constitute reservoirs of functionally relevant antimicrobial resistance determinants shaped by long-term evolutionary processes. In an era of accelerating climate change and increasing ecological connectivity, systematic surveillance of environmental resistance genes will be essential to anticipate their potential dissemination and clinical emergence within a One Health framework.

## METHODS

### Antarctic *Pseudomonas* strains used in this study

Eight Antarctic *Pseudomonas* strains were analyzed in this study. These isolates were recovered from sites located on the Antarctic Peninsula and Union Glacier. Four strains originated from the Henryk Arctowski Polish Station on King George Island, and two were isolated from Union Glacier during the 58th Chilean Scientific Antarctic Expedition (ECA58) conducted in November 2021 (Arros et al., 2024). The remaining strains were obtained from soil samples collected at Coppermine Peninsula (Robert Island) and Yelcho Base (Doumer Island) during the 53rd and 55th Chilean Scientific Antarctic Expeditions carried out between January and February 2017 and 2019 (Marcoleta et al., 2022). Detailed information for each strain is provided in Table S1.

### Cloning and expression of β-lactamases in recombinant strains

AmpC-coding genes from strains PAO1, CoP6a, YeP6b, EH123, and RD2 were cloned into the IPTG-inducible expression vector pET-28b (5368 bp), which confers kanamycin resistance. Recombinant plasmids were transformed into chemically competent *Escherichia coli* BL21 cells prepared using CaCl_2_. Briefly, 5 μL of the plasmid was added to 100 μL of thawed competent cells, incubated at 42°C for 30 seconds, and immediately returned to ice for 5 minutes. Cells were then recovered in 900 µL of LB medium and incubated with shaking for 1 hour at 37°C. After recovery, cells were centrifuged at 7000 rpm for 5 minutes, the supernatant was removed, and 100 µL of the resuspended pellet was plated onto LB agar containing kanamycin. Plates were incubated overnight at 37 °C.

Positive transformants were confirmed by plasmid extraction from overnight liquid cultures using the E.Z.N.A. Plasmid DNA Mini Kit II (Omega Bio-tek, catalog no. D6945-01). Plasmid integrity was verified by restriction digestion with *Xho*I (New England Biolabs) followed by agarose gel electrophoresis.

### Antibiotic susceptibility testing

Disk diffusion assays were performed according to EUCAST guidelines (Matuschek et al., 2014). Overnight cultures of Antarctic strains were grown in Mueller-Hinton (MH) medium at room temperature with rotation, whereas *P. aeruginosa* PAO1 and *E. coli* BL21 were incubated at 37°C with shaking. Cultures were adjusted to OD_625_ = 0.1 in MH medium (final volume 2 mL). A sterile swab was immersed in the suspension and used to inoculate MH agar plates by streaking in three directions to ensure uniform lawn formation. Antibiotic disks (Oxoid) were applied using a disk dispenser. Antarctic isolates were incubated at 25°C for approximately 48 hours, while *E. coli* and *P. aeruginosa* PAO1 were incubated at 37°C for 24 hours.

The following antibiotic disks were tested: cefazolin (CFZ, 30 µg), cefadroxil (CFR, 30 µg), cefoxitin (FOX, 30 µg), cefaclor (CEC, 30 µg), cefotaxime (CTX, 30 µg), ceftazidime (CAZ, 30 µg), ceftolozane-tazobactam (C/T, 40 µg), ceftazidime-avibactam (CAZ-AVI, 50 µg), cefepime (FEP, 30 µg), ampicillin-sulbactam (AMP-SUL, 20 µg), amoxicillin-clavulanic acid (AMC, 30 µg), piperacillin (PIP, 100 µg), piperacillin-tazobactam (TZP, 110 µg), oxacillin (OXA, 1 µg), meropenem (MEM, 10 µg), ertapenem (ETP, 10 µg), and aztreonam (ATM, 30 µg).

Broth microdilution assays were performed following the methodology described by Kadeřábková et al. (2024). The following antibiotics were tested: ceftazidime (maximum concentrations: 256 µg/mL for highly resistant Antarctic strains, 64 µg/mL for less resistant strains, and 4 µg/mL for transformants), cefotaxime (256 µg/mL for highly resistant Antarctic strains, 64 µg/mL for less resistant strains, and 4 µg/mL for transformants), cefepime (8 µg/mL), meropenem (8 µg/mL for Antarctic strains and 0.4 µg/mL for transformants), and ampicillin (256 µg/mL for Antarctic strains and 64 µg/mL for transformants). Each antibiotic was also tested in combination with avibactam (320 µM per well). For transformant strains, IPTG was added at 25 µM to induce gene expression.

### Identification of β-lactam resistance genes from Antarctic genome strains

β-lactam resistance genes were identified using the Resistance Gene Identifier (RGI) tool from the Comprehensive Antibiotic Resistance Database (CARD) (Alcock et al., 2023). Hits were filtered under the drug class category “β-lactam,” retaining sequences with >40% amino acid identity and >80% coverage. To reduce false positives, candidate sequences were subsequently queried against the UniProt database (The UniProt Consortium et al., 2021), and non-antibiotic-resistance-related proteins were excluded.

### Molecular characterization of class C β-lactamases

Class C β-lactamase sequences were aligned using CLUSTAL W (Thompson et al., 1994) and visualized with ESPript (Gouet et al., 1999) to identify conserved catalytic motifs. N-terminal signal peptides were predicted and removed using SignalP (Teufel et al., 2022). Structural alignment-based numbering (SANC) was applied using a hidden Markov model as described by Mack et al. (2020), aligning Antarctic sequences to the Enterobacter cloacae complex P99 reference sequence. A total of 83 residues characteristic of class C β-lactamases were analyzed according to the classification proposed by Philippon et al. (2022).

### Distribution of class C β-lactamases in the P. fluorescens complex

A total of 2,319 Pseudomonas genomes were retrieved from the NCBI RefSeq database (O’Leary et al., 2016) and combined with 27 Antarctic genomes from our laboratory collection, yielding a dataset of 2,346 genomes, including 120 Antarctic Pseudomonas spp. genomes. Taxonomic assignments were validated using GTDB-Tk (Chaumeil et al., 2022), and a phylogenetic tree was constructed with Mashtree (Katz et al., 2019). To further assess genomic relatedness, pairwise average nucleotide identity (ANI) values were calculated using skani (Shaw & Yu, 2023).

Based on ANI comparisons with representative species of the *P. fluorescens* complex (Figure S1) and their clustering within monophyletic clades in the phylogenetic tree, 737 genomes were classified as members of this lineage, including 104 Antarctic genomes. Within this subset, 100 class C β-lactamase sequences were identified using BLASTp (Altschul et al., 1990), applying a minimum protein length of 300 amino acids and an e-value threshold of 1 × 10^−100^.

To further characterize sequence diversity, β-lactamase amino acid sequences were clustered at 90% identity using MMseqs2 (Steinegger & Söding, 2017), incorporating the PDC-1 β-lactamase from Pseudomonas aeruginosa PAO1 as a reference (GenBank ID: AY083595). This reference was included because it represents the best-characterized class C β-lactamase within the genus. Clustering yielded 36 representative sequences, which were used to construct a phylogenetic tree with the NGPhylogeny FastTree pipeline (Lemoine et al., 2019).

