## Supplementary Material for "Intrinsic class C extended-spectrum β-lactamases mediate resistance to oxyimino-cephalosporins in Antarctic *Pseudomonas fluorescens* complex bacteria"

### SUPPLEMENTARY INFORMATION

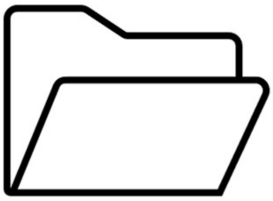

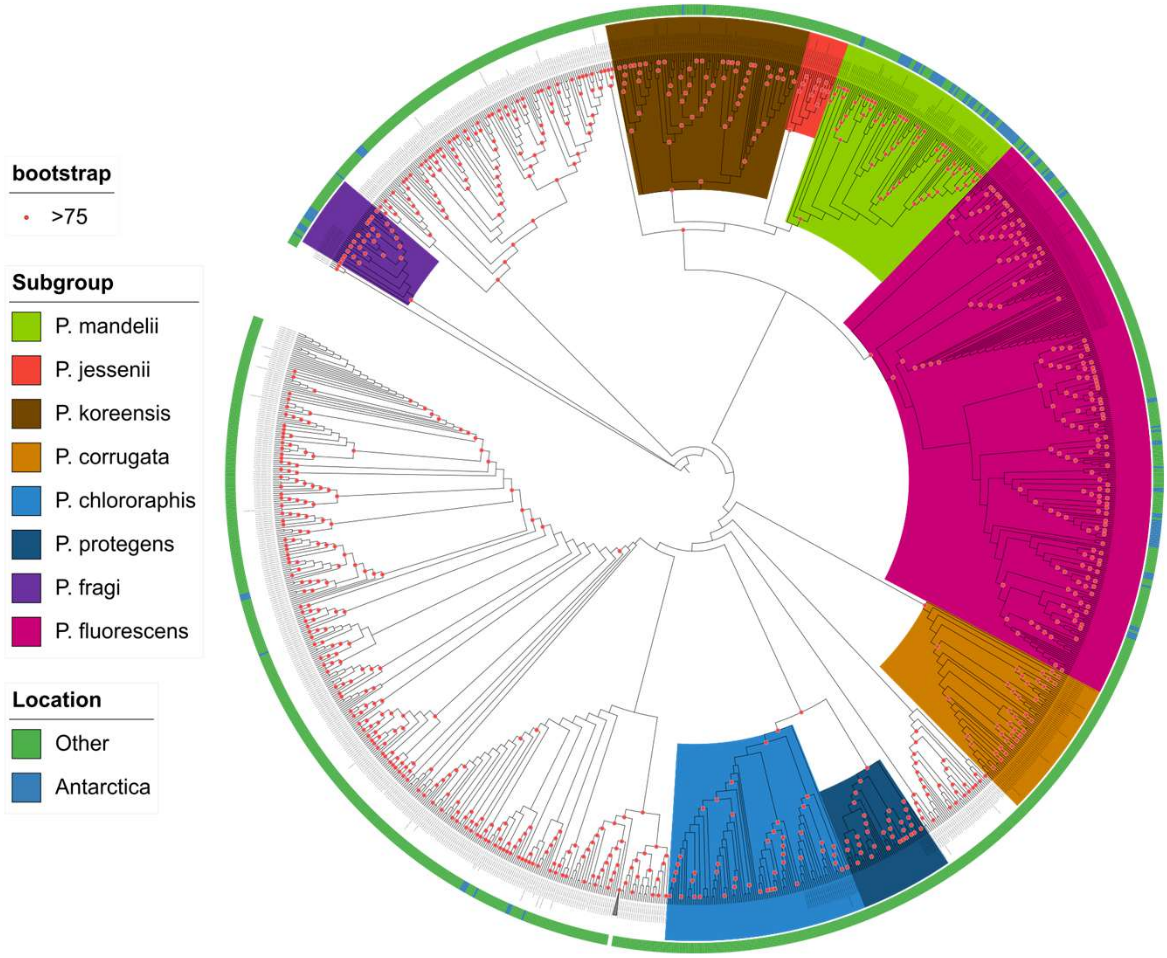


**Figure S1. Phylogenomic structure of the genus *Pseudomonas* and placement of Antarctic *P. fluorescens* complex genomes.** Phylogenomic tree inferred from Mash distances for 2,346 *Pseudomonas* genomes and visualized using iTOL. Colored clades correspond to subgroups within the *Pseudomonas fluorescens* species complex, while uncolored clades represent other *Pseudomonas* lineages, including a collapsed *P. aeruginosa* clade and *P. pertucinogena*. The outermost track denotes geographic origin, with Antarctic isolates highlighted in blue.

**Table S1.** Distribution of class C (AmpC-type) β-lactamases in Antarctic *Pseudomonas fluorescens* complex species and *Pseudomonas aeruginosa*.

| Lineage | Subgroup | N° of sequences | % identity |
|---|---|---|---|
| *P. fluorescens* | all strains | 100 | 48 – 98 |
| | *P. mandelii* | 46 | 58 – 97 |
| | *P. koreensis* | 2 | 94 |
| | *P. fragi* | 10 | 66 – 97 |
| | *P. fluorescens* | 42 | 67 – 98 |
| *P. aeruginosa*[1] | *P. aeruginosa* | 430 | 87-100 |

[1]Values obtained from Philippon et al., 2022 (https://doi.org/10.1128/cmr.00150-21)

**Table S2.** Representative Antarctic class C β-lactamase sequences obtained after clustering at 90% amino acid identity, including the PDC-1 reference enzyme from *Pseudomonas aeruginosa*.

| ***Pseudomonas* species** | **Number of β-lactamase sequences** |
|---|---|
| *P. aeruginosa* | 1 |
| *P. fluorescens* | 14 |
| *P. fragi* | 6 |
| *P. koreensis* | 1 |
| *P. mandelii* | 14 |
| **Total** | 36 |

**Table S3.** Conservation of catalytic residues in representative Antarctic class C β-lactamases based on the SANC numbering system.

| Category | N° analyzed residues | Conservation range (%) |
|---|---|---|
| **Catalytic residues** | 8 | 94.44-100 |
| **Strictly conserved residues (100%)** | 48 | 94-100 |
| **Highly conserved residues (90-97%)** | 34 | 76-100 |

**Table S4.** Geographic origin, isolation source, and taxonomic assignment of Antarctic *Pseudomonas fluorescens* complex strains selected for phenotypic analyses.

| Expedition Year | Antarctic expedition | Strain | Zone | Site | Taxonomy (GTDB-tk) | Subgroup |
|---|---|---|---|---|---|---|
| 2021 | ECA58 | EH123 | Union Glacier | Edson Hills | *Pseudomonas_E frederiksbergensis_F* | *P. mandelii* |
| 2021 | ECA58 | RD2 | Union Glacier | Rossman Cove | *Pseudomonas_E mandelii* | *P. mandelii* |
| 2019 | ECA56 | CoP6a | Robert Island | Coppermine Peninsula | *Pseudomonas_E_sp012935695* | *P. fluorescens* |
| 2019 | ECA56 | YeP6b | Doumer Island | Yelcho Base | *Pseudomonas_E sp002263605* | *P. fluorescens* |
| 2021 | ECA58 | R92 | King George Island | Henryk Arctowski Polish Station | *Pseudomonas_E sp004136015* | *P. mandelii* |
| 2021 | ECA58 | S135 | King George Island | Henryk Arctowski Polish Station | *Pseudomonas_E sp004136015* | *P. mandelii* |
| 2021 | ECA58 | S137 | King George Island | Henryk Arctowski Polish Station | *Pseudomonas_E sp004136015* | *P. mandelii* |
| 2021 | ECA58 | S134 | King George Island | Henryk Arctowski Polish Station | *Pseudomonas_E sp900187425* | *P. mandelii* |

**Table S5.** Amino acid sequence identity between Antarctic class C β-lactamases and closest homologs in the curated β-lactamase databases BLDB and CARD.

| Gene | ***bla*$_{PAO1}$ (PDC-1)** | **CARD best hit (%)** | **BLDB best hit (%)** |
|---|---|---|---|
| *bla*$_{CoP6a}$ | 62.60 | PDC-362 (56.04) | PFL-5 (83.59) |
| *bla*$_{EH123}$ | 61.90 | PDC-455 (60.89) | PFL-6 (66.67) |
| *bla*$_{RD2}$ | 61.30 | PDC-455 (60.37) | PFL-4 (62.95) |
| *bla*$_{YeP6b}$ | 62.10 | PDC-241 (58.36) | PFL-4 (73.33) |

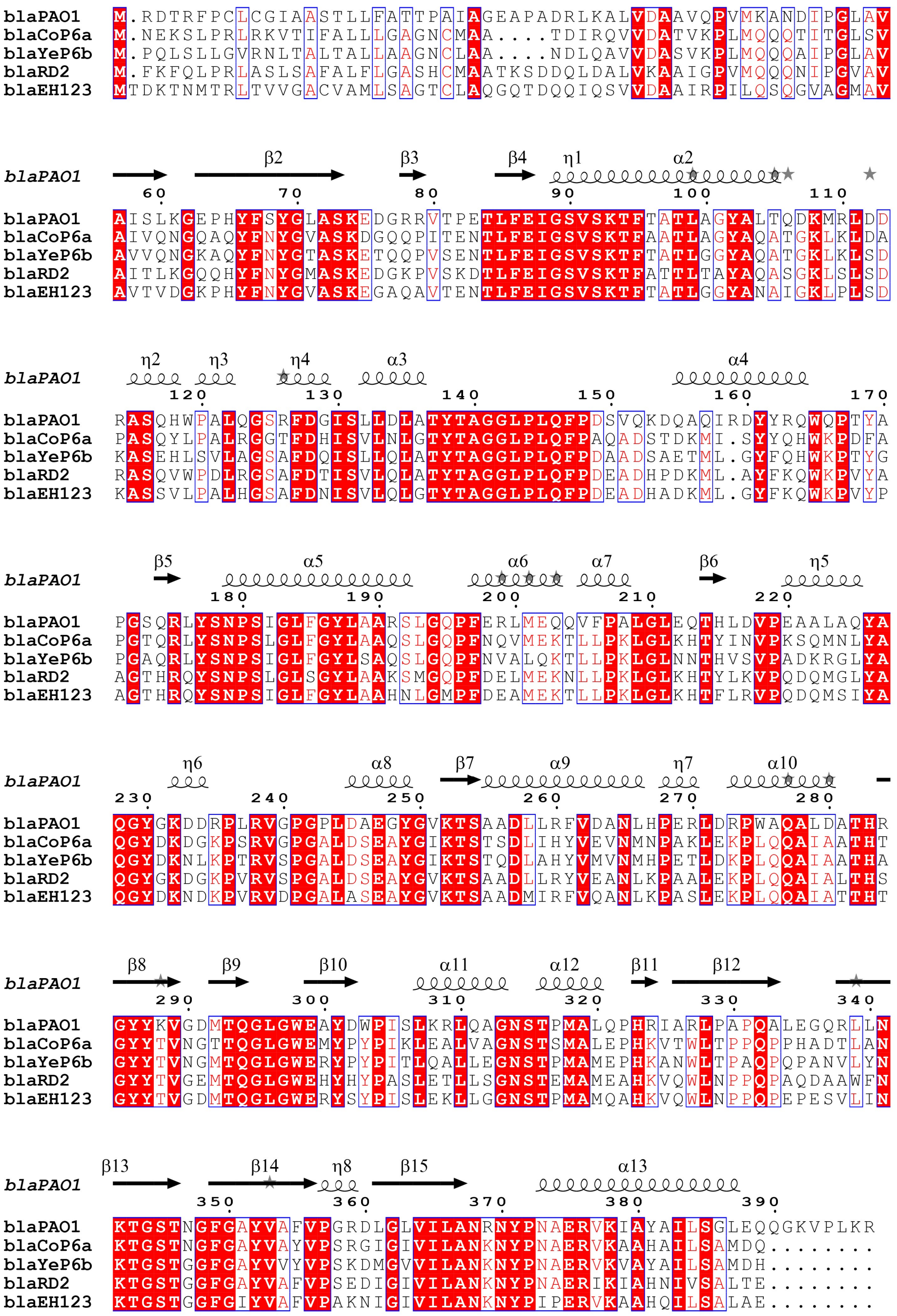


**Figure S2. Conservation of catalytic motifs in Antarctic class C β-lactamases.** Multiple amino acid sequence alignment of representative Antarctic class C β-lactamases and the PDC-1 reference enzyme from *P. aeruginosa* PAO1. Canonical class C catalytic motifs (S64XSK, Y150XN, and K315TG) are indicated by black boxes. Residue numbering follows the structural alignment–based numbering (SANC) system.

**Table S6.** Analysis of strictly and highly conserved residues in selected Antarctic class C β-lactamases and the PDC-1 reference enzyme.

| | **Conserved residue category of class C beta-lactamase** | | |
|---|---|---|---|
| **β-lactamase** | **Catalytic** | **Strictly conserved** | **Highly conserved** |
| *bla*$_{PAO1}$ (PDC-1) | 8/8 (100%) | 47/48 (98%) | 34/34 (100%) |
| *bla*$_{CoP6a}$ | 8/8 (100%) | 47/48 (98%) | 29/34 (85%) |
| *bla*$_{EH123}$ | 8/8 (100%) | 48/48 (98%) | 33/34 (97%) |
| *bla*$_{RD2}$ | 8/8 (100%) | 47/48 (98%) | 33/34 (97%) |
| *bla*$_{YeP6b}$ | 8/8 (100%) | 47/48 (98%) | 32/34 (94%) |

**Table S7.** Minimum inhibitory concentrations of β-lactam antibiotics for recombinant *Escherichia coli* BL21 strains expressing Antarctic class C β-lactamases and PDC-1, in the presence or absence of avibactam.

| ***bla* gene** | **MIC (µg/mL)** | | | | | |
|---|---|---|---|---|---|---|
| | AMP | AMP-AVI | CTX | CTX-AVI | CAZ | CAZ-AVI |
| $bla_{PAO1}$ | 64 | 8 | 4 | <0.125 | 0.5 | <0.125 |
| $bla_{EH123}$ | 32 | <2 | 4 | <0.125 | 4 | <0.125 |
| $bla_{RD2}$ | 64 | <2 | >4 | 0.25 | 4 | <0.125 |
